# A novel small molecule screening platform for disrupting toxic tau oligomers in cells

**DOI:** 10.1101/510412

**Authors:** Chih Hung Lo, Colin Kin-Wye Lim, Zhipeng Ding, Sanjula Wickramasinghe, Anthony R. Braun, Elizabeth Rhoades, David D. Thomas, Jonathan N. Sachs

## Abstract

Tauopathies, including Alzheimer’s disease, are a group of neurodegenerative disorders characterized by pathological aggregation of the microtubule binding protein tau. Recent studies suggest that toxic tau oligomers, which are soluble and distinct from insoluble beta-sheet fibrils, are central players in neuronal cell death. To exploit this new therapeutic window, we engineered two first-in-class FRET based biosensors that monitor tau conformations in cells. Because this new technology platform operates in cells, it enables high-throughput screening of small molecules that target tau oligomers while avoiding the uncertainties of idiosyncratic *in vitro* preparations of tau assemblies from purified protein. We found a small molecule, MK-886, that disrupts tau oligomers and reduces tau-induced cell cytotoxicity with nanomolar potency. Using SPR and an advanced single-molecule FRET technique, we show that MK-886 directly binds to tau and specifically perturbs the folding of tau monomer in the proline-rich and microtubule-binding regions. Furthermore, we show that MK-886 accelerates the tau aggregation lag phase using a thioflavin-T assay, implying that the compound stabilizes a non-toxic, on-pathway oligomer. The technology described here should generalize to the study and targeting of conformational ensembles within the aggregation pathways of most intrinsically disordered proteins.

## Introduction

Tauopathies, including Alzheimer’s disease (AD), are a group of neurodegenerative disorders characterized by the presence of tau inclusions (neurofibrillary tangles, NFTs) in affected brain regions^1^. Despite the past few decades of rigorous and focused research, there are currently no cures or significant disease modifying therapies for AD and related tauopathies^2^. There is a particular dearth of compounds that target tau, as out of the 105 small molecules that are currently in clinical trials for AD, only five are tau-related disease modifying small molecules^3–4^. Hence, there is desperate need for technologies that enable the identification of more effective disease-modifying and tau-focused treatments^4–6^.

Tau is an intrinsically disordered protein that plays an important role in the regulation of microtubule stability and axonal transport^7^. Under pathological conditions, tau is hyperphosphorylated and detaches from microtubules, accumulating in the cytosol^8^. Unbound tau has a tendency to misfold, undergoing conformational changes that initiate the tau amyloidgenesis cascade, with an initial formation of tau oligomers that subsequently nucleate into paired helical filaments (PHFs), and eventually intracellular NFTs^9^. NFTs have been the primary histopathological hallmark of tauopathies, with their presence in the brain showing significant correlation with the degree of cognitive impairment^10^. However, recent studies suggest that these large insoluble NFTs are not the principle toxic species, implicating soluble oligomeric tau— intermediate tau assemblies formed prior to PHFs and NFTs—in the induction of neurodegeneration^11–12^. Tau oligomers promote cytotoxicity *in vitro* and are linked to neurodegeneration and cognitive phenotypes *in* vivo^12–18^. As a result, the focus in therapeutic development has begun to shift from targeting large fibrillar aggregates to inhibiting or disrupting the formation of toxic soluble tau oligomers^11, 19–21^.

Several recent efforts to discover small molecules that target toxic tau oligomers have yielded efficacious, cytoprotective compounds^22–28^. However, these studies were done *in vitro* with purified proteins, and the compounds were protective only in the low micromolar range^22–28^. Tau oligomers exist as an ensemble of distinct assemblies which include both toxic and non-toxic, on- and off-pathway species along the fibrillogenesis cascade^29–35^. The formation of these toxic tau oligomers has been associated with mutations and overexpression of numerous chaperone proteins^36–37^, highlighting the importance of other components of the tau-protein interactome in the pathogenesis of tauopathies. Capturing this complexity in an *in vitro* setting is extremely difficult, if even possible, and established protocols for aggregating purified tau protein into oligomers, PHFs, and NFTs have, not surprisingly, been shown to produce different tau assemblies depending on aggregation conditions^17, 31, 34, 38–41^. Critically, no specific toxic tau species has been isolated or identified to date^21, 42–43^.

Here, we shift the tau oligomerization process into the cellular environment, where the ensemble of tau conformations should more closely recapitulate the distribution of oligomeric assemblies. In addition, cellular oligomers, unlike those produced from purified proteins, may include non-tau components and thus tau oligomer structures only accessible via interaction with chaperone proteins. Building on the groundbreaking biosensor developed by the Diamond group (which is focused on the detection of pathogenic species in biofluids as a biomarker for AD diagnosis)^44^, we have developed a technology platform that directly monitors tau oligomerization in cells, enabling the therapeutic targeting of early-stage tau pathology. Furthermore, with this platform, we have established a robust assay that can be easily adjusted in the future for additional cell lines and new pathological conditions that more closely mimic disease conditions as they become known. Such a platform increases the likelihood of targeting the true toxic species, and has the added advantage of identifying compounds that act both directly (by binding tau) and indirectly (through orthogonal biochemical pathways) to modify toxic oligomers.

To discover small molecules that disrupt toxic tau oligomers in cells, we engineered two distinct fluorescence resonance energy transfer (FRET) biosensors to monitor tau oligomerization. We used these biosensors in conjunction with a state-of-the-art fluorescence lifetime plate reader as a high-throughput screening (HTS) platform for drug discovery^45^, using the NIH Clinical Collection (NCC) library. Fluorescence lifetime detection increases the precision of FRET-based screening by a factor of 30 compared with conventional fluorescence intensity detection^46^, and provides exquisite sensitivity to resolve minute structural changes within protein ensembles. This sensitivity allows direct detection of conformation changes within an ensemble of oligomers (e.g. conversion from toxic to non-toxic oligomer conformation), the dissociation of oligomers, or even changes in the ensemble of monomer conformations^47–49^. The FRET biosensors were engineered by expressing full-length wild-type (WT) 2N4R tau and fluorescent protein fusion constructs in living cells, allowing us to directly detect and monitor either *inter-molecular* or intra-molecular tau interactions. While FRET techniques have been employed to investigate tau aggregation with aggregation-prone constructs, such as the caspase-cleaved form of tau^50^ or the K18 isoform with familial mutants including P301L^44^, the use of full-length WT 2N4R tau to study tau oligomerization has not been reported. The novel use of full-length WT 2N4R tau in this study enables the specific detection of ensembles of tau oligomers, not fibrils, as the 2N4R isoform does not spontaneously fibrillize, even at supersaturating concentrations, and therefore forms mostly oligomers^51–53^.

After first establishing that the new technology platform specifically targets oligomers (using known tau aggregators), we identified a small molecule, MK-886, that directly binds tau and strongly attenuates FRET with an EC50 of 1.40 μM. The compound rescues tau-induced cell cytotoxicity with an IC50 of 435 nM. To elucidate the mechanism of action, we used an advanced single-molecule FRET (smFRET) technique to show that MK-886 perturbs the folding of purified, monomeric tau in the proline-rich and microtubule-binding regions. This effect is recapitulated in the cellular biosensor that monitors intra-molecular FRET and indicates an unfolding of the two termini of tau. In addition, we used a thioflavin-T assay to show that MK-886 shortens the lag phase in tau fibrillization, implying that the compound stabilizes a non-toxic, on-pathway tau oligomer, hence defining a new therapeutic approach to targeting toxic tau oligomers.

## Results

### Inter-molecular FRET biosensor directly monitors structural changes in tau oligomers in cells

To develop an in-cell HTS platform that can detect small-molecule modulation of tau oligomerization and/or perturbation of tau conformational states, we engineered a tau FRET biosensor expressed in living cells. We used HEK293 cells expressing full-length WT 2N4R tau fused to GFP or RFP (tau-GFP/RFP or tau FRET biosensor) (**Fig. 1A**). Full-length WT 2N4R isoform was used because it does not spontaneously fibrillize, even at supersaturating concentrations, forming mostly oligomers^51–53^. Expression and homogeneity of the FRET biosensor were determined by fluorescence microscopy and immunoblotting. Fluorescence microscopy images showed that the tau proteins were evenly distributed in the cytosol of the cells, with no discernable puncta (which would have indicated more progressive aggregation, e.g. fibril formation) or other non-uniformities (**Fig. 1B**). Western blot analysis of the tau biosensor cell lysates confirmed the expression of fluorophore-tagged tau (**Supplementary Fig. 1A**).

**Fig. 1:**
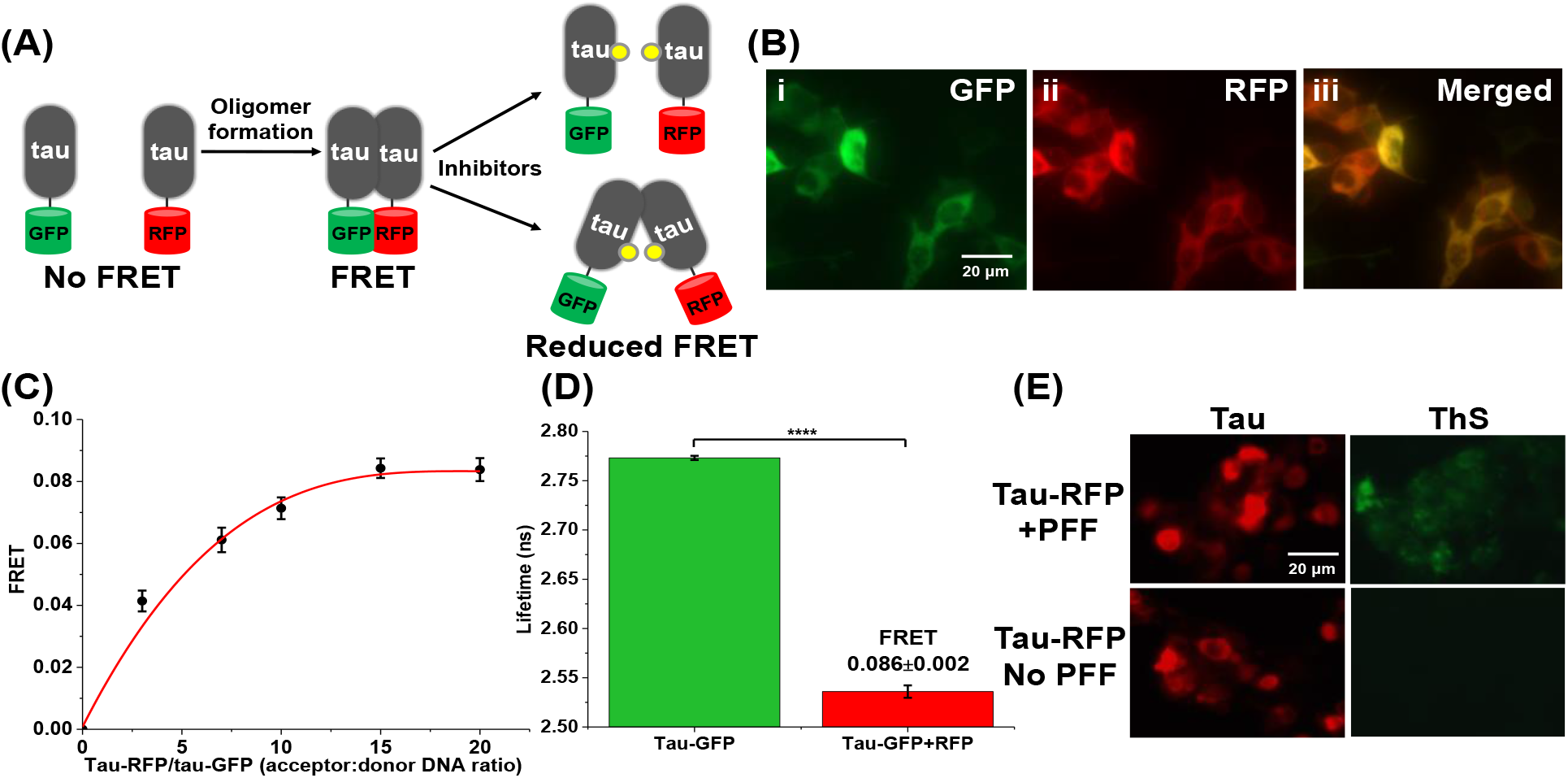
The tau *inter*-molecular FRET biosensor and fluorescence lifetime technology enables direct monitoring of tau oligomerization in cells. (A) Schematic representation of live-cell based tau *inter*-molecular FRET biosensor. FRET signal is observed when tau oligomers form, which is modulated by small-molecule inhibitors. Tau oligomer is drawn as a dimer for illustration but it can be more than a dimer (≥2-mers). (B) Fluorescence microscopy images of HEK293 cells expressing tau-GFP/RFP (tau FRET biosensor). (i) GFP channel, (ii) RFP channel, (iii) merged channel showing the presence of both tau-GFP and tau-RFP colocalized in the cells. (C) FRET efficiency between tau-GFP (donor) to tau-RFP (acceptor) shows hyperbolic dependence on acceptor concentration. (D) The mean fluorescence lifetime of donor-only control is τ_D_=2.773±0.002 ns and in the presence of saturating acceptor is τ_DA_=2.536±0.006 ns, corresponding to a FRET efficiency E=0.086±0.002, indicating tau self-association. (E) Thioflavin-S (ThS) staining of HEK293 cells expressing tau-RFP (in same total DNA concentration as used in biosensor) confirms the absence of β-sheet tau species. Only cells treated with preformed tau fibrils (PFF) showed a positive ThS signal while untreated cells expressing tau-RFP only were not ThS positive. N=3 and ****p<0.0001.

We next tested the functionality of the tau FRET biosensor by measuring FRET efficiency using the fluorescence lifetime plate reader^45^. FRET between tau-GFP (donor) and tau-RFP (acceptor) in live cells showed hyperbolic dependence on acceptor concentration (**Fig. 1C**), with a maximum energy transfer efficiency (E) of 0.086±0.002, illustrated through a substantial decrease in the donor fluorescence lifetime in the presence of the acceptor (**Fig. 1D**), indicating the formation of tau oligomers in cells. As a negative control, we expressed free soluble GFP and RFP in cells using the same DNA concentrations as the tau biosensor to ensure that the FRET we observed was tau-specific and not caused by nonspecific interactions between the fluorophores in the cytosol (**Supplementary Fig. 1B**). The low FRET efficiency (E=0.019±0.004 between free soluble GFP and RFP) indicates that FRET observed from cells expressing tau biosensor arises from specific tau-tau interactions and not from nonspecific interactions between the free fluorophores (**Supplementary Fig. 1C**).

We further tested the sensitivity of the tau FRET biosensor with the addition of forskolin, a small molecule known to induce tau hyperphosphorylation and self-association^54^. Cell treatment with forskolin (20 μM) showed a significant increase in FRET, illustrating increased self-association of tau (**Supplementary Fig. 1D**). Interestingly, cell treatment with gossypetin (10 μM), a known inhibitor of fibril formation^27^, did not show any significant change in FRET, suggesting that under our cellular conditions only oligomeric tau species were contributing to the observed FRET (**Supplementary Fig. 1D**).

To further confirm that only oligomeric species of tau, but not fibrils, were present in the tau biosensor cells, we performed a thioflavin-S (ThS) assay in cells expressing tau-RFP at the same concentration of tau-GFP/RFP dual transfected cells, with treatment of tau pre-formed fibrils (PFF) as positive controls. Tau-RFP was used instead of tau-GFP/RFP as tau-GFP emission interferes with the ThS signal. Results from the ThS assay illustrate that the cells treated with PFF show a positive ThS signal, but not the cells expressing the biosensors (**Fig. 1E**), confirming that no fibrils (specifically β-sheet tau assemblies) are present in the biosensor cells, and more importantly that the observed FRET is mainly the result of tau oligomerization. The combination of our biosensor’s basal FRET signal and response to positive control tool compounds demonstrates that time-resolved FRET detection in cellular tau FRET biosensors is sensitive to conformationally distinct tau assemblies, providing a powerful platform to identify novel compounds that modulate the ensemble of tau oligomers.

### Identification of novel small molecules from HTS of the NCC library that perturb the conformational ensembles of tau oligomers

Using our cellular tau FRET biosensor, we performed a HTS of the NIH Clinical Collection (NCC: 727 bioactive compounds) to identify compounds that perturb the conformational ensembles of tau oligomers. The NCC library is a collection of small molecules that have been previously tested in clinical trials, and therefore have known safety profiles and details on potential mechanisms of action. These compounds can provide excellent starting points for medicinal chemistry optimization and may even be appropriate for direct human use in new disease areas.

After an initial quality control check of the cells expressing the tau FRET biosensor on each day of screening (fluorescent waveform signal level and coefficient of variance), the cells were dispensed into drug plates and incubated with the compounds (10 μM) or DMSO control wells for 2 hours. Lifetime measurements were acquired with the fluorescence lifetime plate reader. A single-exponential fit was used to determine the lifetime from cells expressing the tau FRET biosensor (γda) or expressing a tau-GFP donor-only control (γd) to determine FRET efficiency (Eq. 1). As fluorescence lifetime measurements are prone to interference from fluorescent compounds, a stringent fluorescent compound filter was used to flag 30 such fluorescent compounds as potential false-positives due to interference from compound fluorescence^47–48^. FRET efficiency from all compounds that passed the fluorescent compound filter are plotted (**Fig. 2A**) and a histogram of the FRET distribution from these compounds was fit to a Gaussian curve to obtain a mean and standard deviation (σ, SD) for the screen (**Supplementary Fig. 2A**).

**Fig. 2:**
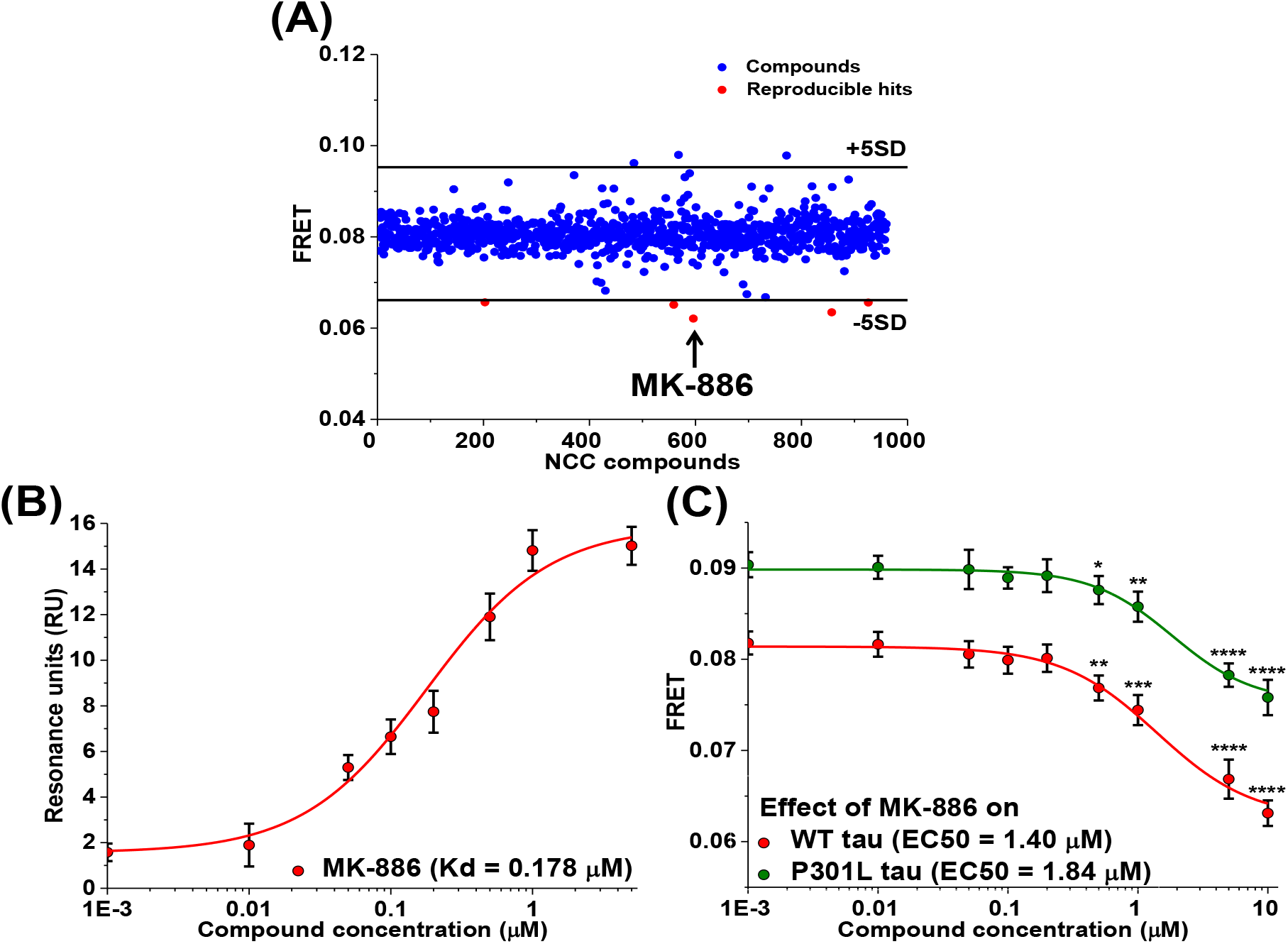
Identification of MK-886 as a compound that directly perturbs conformational ensemble of tau oligomers from high-throughput screening of NIH clinical collection (NCC) library. (A) Representative pilot screening with NCC library containing 727 compounds. A FRET efficiency cutoff threshold was applied at a change in FRET efficiency of 0.015 (5SD, *black lines).* Five reproducible hits that decrease FRET by more than 5SD below the mean of all cells were identified from the pilot screens (red). (B) Surface plasmon resonance (SPR) binding curve of MK-886 to purified recombinant full-length wild-type (WT) 2N4R tau proteins shows that the binding affinity of MK-886 was 0.178 μM. MK-886 also has the largest FRET change (arrow in (A)). (C) FRET dose-response assays with MK-886. The compound produced a dose-dependent FRET change in both WT and P301L *inter*-molecular biosensors with EC50 of 1.40 and 1. 84 μM respectively. N=3, *p<0.05, **p<0.01, ***p<0.001 and ****p<0.0001.

Our initial goal was to discover compounds that alter the conformational ensembles of tau oligomers with the potential of disrupting tau-tau interactions, leading us to focus our search to compounds that reduce FRET (though other compounds that increase FRET could potentially remodel toxic oligomers and be of interest in future studies). Five reproducible hits from the library were shown to decrease FRET by more than 5σ below the mean of all wells while not appearing as hits in the donor-only control screen (**Fig. 2A**, highlighted in red and **Supplementary Fig. 2B-C**).

### Binding of hit compounds to purified tau proteins

In the present study, we are focused on identifying compound that directly interact with tau and modulate the FRET signal from tau oligomers. To determine if these five hit compounds directly act on tau or modulate tau FRET by acting through an indirect pathway, we measured the binding affinity for each of the five hit compounds to purified tau using surface plasmon resonance (SPR). SPR measurements were carried out by immobilizing purified recombinant full-length WT 2N4R tau onto the SPR chip followed by compound flow through the chip to allow for binding. Of the five hits that reduced FRET with our tau biosensor, MK-886 was the only hit to demonstrate dose-dependent binding to purified tau protein; the measured binding affinity (Kd) is 178 nM (**Fig. 2B** and **Supplementary Fig. 2D**). Interestingly, MK-886 had the strongest change in FRET (**Fig. 2A**, highlighted in arrow). The other four hit compounds did not show direct binding to immobilized tau protein (**Supplementary Fig. 2E**) and therefore most likely attenuate the tau oligomer FRET signal via an indirect mechanism of action. All subsequent analysis in this study is focused on the direct tau binder, MK-866, however the indirect effectors of tau oligomerization are potentially useful and are briefly discussed below.

### FRET dose-response of MK-886 with cellular tau *inter*-molecular FRET biosensors

The relative effective concentration (EC50) of MK-886 was determined by in-cell FRET measurements using the wild-type tau biosensor. The compound decreased FRET efficiency in a dose-dependent manner (**Fig. 2C**), with EC50 of 1.40 μM. To test the effect of MK-886 on a more disease-relevant model, we also performed a FRET dose response experiment using a P301L tau FRET biosensor, a more aggregation prone mutant of tau^37^. The FRET efficiency of this biosensor was found to be 0.091, which is higher than that of WT tau biosensor, consistent with the known tendency of P301L tau to be hyperphosphorylated and hence more oligomeric^37^. Similar to the wild-type, we observed a dose-dependent decrease in the FRET efficiency of the P301L tau biosensor with MK-886 with an EC50 of 1.84 μM (**Fig. 2C**), confirming that the hit compound remodels tau oligomers in a disease-relevant model.

To ensure that MK-886 was not reducing FRET through interactions with the fluorophores, we performed controls with cells expressing GFP-linker-RFP (linker contains 32 amino acids, GFP-32AA-RFP) (**Supplementary Fig. 3A**). FRET measurements with this GFP-32AA-RFP (**Supplementary Fig. 3B**) as well as soluble GFP and RFP (**Supplementary Fig. 3C**) showed no change in FRET efficiency in the presence and absence of MK-886, which confirmed that the small molecule was not acting directly on the cytosolic fluorophores. Assay quality *(Z’)* was determined using MK-886 (Eq. 2). The *Z’* value of 0.72±0.02 indicates excellent assay quality, validating MK-886 as a positive control tool-compound for targeting tau oligomers.

### MK-886 reduces tau-induced cell cytotoxicity in SH-SY5Y cells with nanomolar potency

We next tested the effect of MK-886 on tau-induced cytotoxicity in the SH-SY5Y neuroblastoma cell model of tauopathy^17, 22, 55^. Overexpression of P301L tau showed significantly greater cell death (30%) when compared to the vector control (18%) (**Fig. 3A**). Treatment with MK-886 (1 nM to 5 μM) to cells overexpressing P301L tau showed significant, though incomplete, rescue of tau-induced cytotoxicity in a dose-dependent manner, with an IC50 of 435 nM (**Fig. 3B**), the same order of magnitude as MK-886’s binding affinity for recombinant tau protein. We note that the IC50 of MK-886 in the cell cytotoxicity assay was lower than the EC50 observed from the FRET assay. This is likely due to the different conditions optimized for each assay, specifically incubation time and cell type (72 hours incubation in SH-SY5Y cells for the cytotoxicity assay versus 2 hours incubation in HEK293 cells for the FRET assay).

**Fig. 3:**
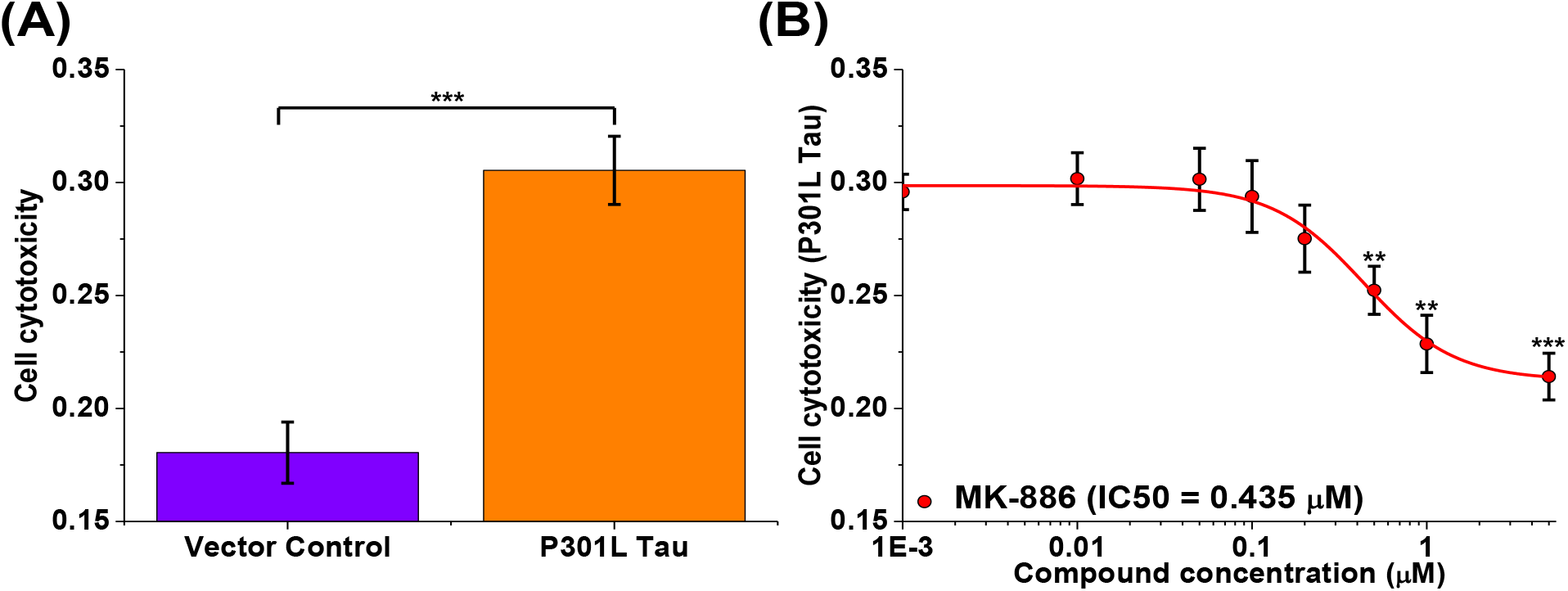
Inhibition of tau-induced cell cytotoxicity in SH-SY5Y human neuroblastoma cell line by MK-886. (A) SH-SY5Y human neuroblastoma cells were transfected with vector control and P301L tau. Significant cell death is observed in cells transfected with P301L tau (30%) as compared to the vector control (18%). (B) Hit compound (MK-886) inhibited P301L tau-induced cytotoxicity in SH-SY5Y with an IC50 of 0.435 μM. N=3, **p<0.01 and ***p<0.001.

We note that MK-886—which was blindly identified in our HTS—has been shown to play a role in modulating AD-related amyloid and tau pathology through inhibition of 5-lipoxygenase (5-LOX)-activating protein (FLAP)^56^, potentially altering the clearance and phosphorylation state of tau^57–58^. Our observations suggest that MK-886 rescues tau-induced cytotoxicity through direct binding to tau protein and not by modulating FLAP, a previously undescribed mechanism of action. SH-SY5Y cells do not express 5-LOX or FLAP and therefore are a particularly well suited model to evaluate alternative mechanisms of action for MK-886 rescue of tau-induced cytotoxicity^59^. We also confirmed that there were no changes in the relative levels of expressed tau (**Supplementary Fig. 4A**) or the phosphorylation state (**Supplementary Fig. 4B**) due to MK-886 treatment in the SH-SY5Y cell model. These results support our hypothesis that MK-886 rescues tau-induced cytotoxicity by direct perturbation of the ensemble of conformational states of toxic tau oligomers and provides a novel molecular mechanism through which MK-886 may have an effect on tauopathies.

### MK-886 specifically perturbs the PRR/MTBR of tau monomer and induces conformational changes of the cellular tau intra-molecular biosensor

To further investigate the mechanism of action of MK-886, we used single-molecule FRET (smFRET) to examine the effect of MK-886 on monomeric tau. Using two different doubly fluorescent-labeled tau constructs (labeled at the proline rich region/microtubule binding region (PRR/MTBR) or at the N-terminal domain) as previously developed in our lab^60^, we monitored the conformation of two distinct regions of the tau protein (**Fig. 4A**). The smFRET shows that the addition of MK-886 results in a substantial increase in FRET in the PRR and the MTBR targeted construct (**Fig. 4B**) but only a minor decrease in FRET for the construct monitoring the N-terminal domain conformation (**Fig. 4C**). This suggests that MK-886 specifically binds and induces a conformational change in tau monomer at the PRR/MTBR region (increased FRET), resulting in a subsequent loss of interactions between the N-terminal domain and the PRR/MTBR (decreased FRET) in the second construct. To determine whether MK-886 also perturbs the monomer conformation of tau in cells, we tested the compound with a cellular tau *intra*-molecular FRET biosensor (GFP-tau-RFP) expressed in HEK293 cells. The *intra*-molecular FRET biosensor has a basal 6% FRET signal (**Supplementary Fig. 5**), illustrating the *intra*-molecular interactions arising from the paper-clip monomeric structure in which the N- and C-terminus of tau are folded to close proximity^61^. Treatment with MK-886 reduced the *intra*-molecular FRET with an EC50 of 2.12 μM, similar to that of oligomer modulation, suggesting that the change in conformational states of oligomers is due, are at least correlated with, conformational perturbation of the tau monomer (**Fig. 4D**).

**Fig. 4:**
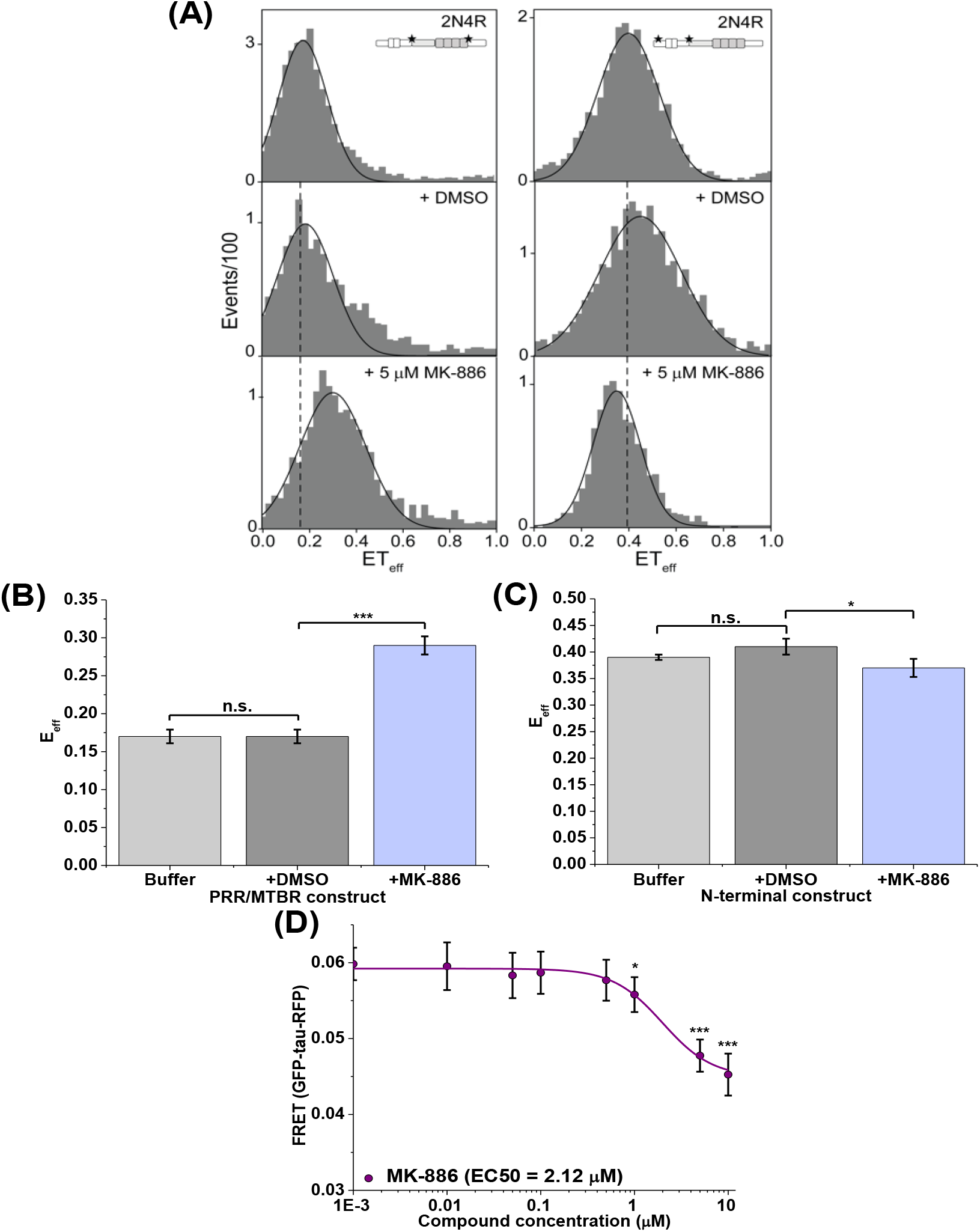
MK-886 binds and perturbs tau monomer conformation. (A) Single-molecule FRET (smFRET) measurements in the absence or presence of MK-886 with full-length wild-type 2N4R tau labelled at the proline-rich region/microtubule binding region (PRR/MTBR) (left) or at the N-terminal domain (right). Tau schematic represents the labelling position for each construct. The black line is drawn from the peak of the histogram in buffer for comparison with DMSO and MK-886 measurements. Histograms shown are representative. (B-C) Quantification of the smFRET measurements from (A) indicates that PRR/MTBR become substantially more compact upon binding MK-886 (5 μM) (A, bottom panel) when compared to tau in buffer (A, top panel) or DMSO (A, middle panel) while the N-terminal domain shows only minor differences in the presence of MK-886 (5 μM). (D) MK-886 also produced a dose-dependent FRET change in the cellular tau *intra*-molecular biosensor with EC50 of 2.12 μM, similar to that of oligomer modulation, suggesting that the change in conformational states of oligomers is due in part to conformational changes of tau monomer. N=3, *p<0.05, ***p<0.001 and n.s. indicates not significant.

### MK-886 stabilizes tau conformations that promote the formation of β-sheet-positive fibrils in the presence of aggregation inducer

We have shown above that MK-886 is capable of directly binding to immobilized recombinant tau protein, modulates tau oligomer and monomer conformation (both in cells and in purified proteins), and rescues tau-induced cytotoxicity. To further investigate the mechanism of action and identify whether MK-886 targets on- or off-pathway oligomers, we performed a heparin-induced thioflavin-T (ThT) aggregation assay in the absence and presence of MK-886. MK-886 shortens the lag phase of tau β-sheet fibril formation in a dose-dependent manner (**Fig. 5A**), suggesting that it induces or stabilizes on-pathway, early species in the amyloidogenic cascade. We confirmed that MK-886 did not have a direct effect on ThT fluorescence (**Supplementary Fig. 6A**) and did not act as a separate nucleation point for fibril formation, as incubation of tau protein with MK-886 in the absence of heparin did not lead to ThT-positive signals (**Supplementary Fig. 6B**). In addition, we tested if MK-886 disrupts tau preformed fibrils (PFF). Comparison of MK-886 to gossypetin (a known remodeler of tau fibrils) illustrates that MK-886 did not reduce the ThT signal from tau PFF, whereas gossypetin showed a significant decrease, indicating the disruption of β-sheet fibril structure (**Fig. 5B**). These results, in combination with the changes in FRET and reduction of tau-induced cytotoxicity, suggest that MK-886 alters the conformational ensemble of tau oligomers favoring a subset of non-toxic, on-pathway oligomers that promote tau fibrillization.

**Fig. 5:**
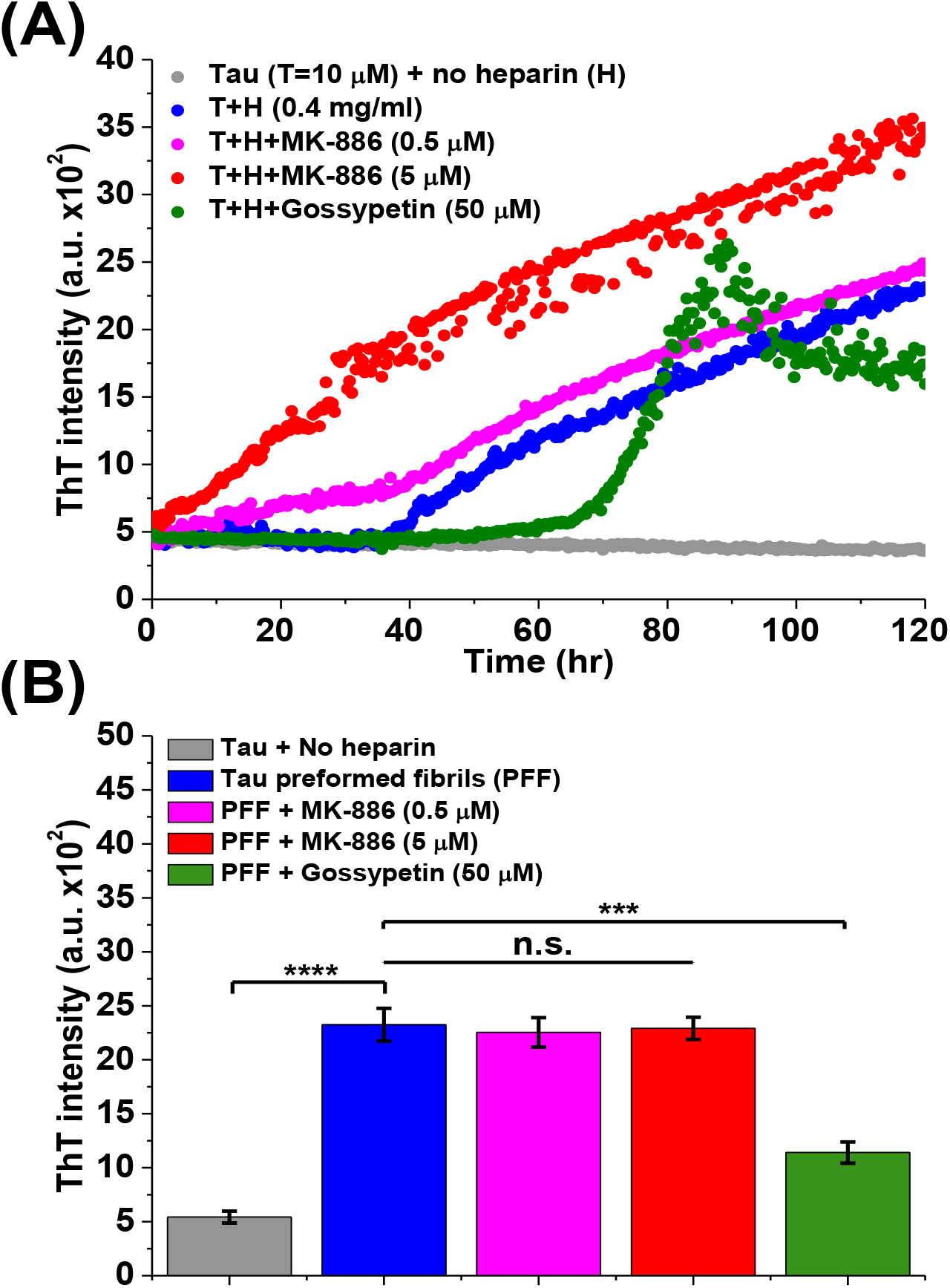
MK-886 alters the ensemble of conformational states of tau oligomers by stabilizing a fibrillization promoting conformation. (A) Thioflavin-T assay was performed with purified recombinant full-length wild-type 2N4R tau. Fibrillization was induced by heparin in the presence of DMSO control (2% v/v), hit compound (MK-886, 0.5 and 5μM) and known a small-molecule inhibitor in remodeling tau fibrils (gossypetin, 50μM). Results indicate that MK-886 reduces a lag phase of β-sheet fibril formation. (B) Incubation with compound MK-886 and gossypetin to preformed fibrils (PFF) for 72 hours shows that MK-886 does not disrupt or remodel PFF while gossypetin disrupts PFF. All samples were treated with DTT (5 mM). N=3, ****p<0.0001 and n.s. indicates not significant.

## Discussion

Although small molecules that inhibit or disrupt tau fibril formation have been studied for decades, the targeting of toxic tau oligomers has only recently been reported, with the majority of efforts being focused on repurposing previously identified small molecules that disrupt tau fibril formation. These small molecules include phenothiazines such as methylene blue (MB) and its derivatives Azure A, Azure B and Azure C (K_d_=100 nM-3.4 μM; fibrillization IC_50_=1-10 μM; Azure C rescues cell cytotoxicity at 5 μM^)22-23, 27^, polyphenols such as epigallocatechin gallate (EGCG) (cell cytotoxicity and fibrillization inhibited at 12.5 μM)^24^ and curcumin (K_d_=3.3 μM; fibrillization IC_50_=10.6 μM)^25^, and heparin-like oligosaccharides (K_d_=140-350 nM; cell cytotoxicity rescued at 10 μM)^26^. Of these small molecules, only MB has reached phase III clinical trials, ultimately with unsuccessful results as no significant improvement of patient cognitive outcomes were reported^62^.

One commonality among these molecules is that they were initially identified through *in vitro* purified protein assays^22–27^, which do not reliably represent the cellular environment. For example, these assays lack numerous chaperone proteins that may contribute to the continuum of toxic and non-toxic, on- and off-pathway tau oligomers that populate the amylogenic cascade. Inherently, these purified protein assays are only capable of identifying hits that directly perturb tau protein and are wholly naive against indirect mechanisms of action, affecting other pathologically relevant cellular processes. Furthermore, many of the small molecules discovered in purified protein assays disrupt not only tau oligomers but also fibrils, which have been suggested to be inert and potentially neuroprotective^63^. Hence, a cellular approach that monitors tau oligomers and not fibrils holds promise as a novel HTS platform to discover more effective therapeutics.

Our small molecule hit compound, MK-886, shows strong binding affinity for recombinant tau (K_d_ = 178 nM, comparable to other known tau aggregation inhibitors) and with improved potency for tau-induced cytotoxicity (IC_50_ = 435 nM). We observed that MK-886 changes the conformation of the PRR and MTBR domains within the tau monomer. It has been suggested that the folding over of the two termini of tau to form the classic “paper-clip” monomeric structure is due to electrostatic interactions that arise from the opposite net charges of the N-terminal and MTBR domains^61^. While this global folding is specific, it has been shown to be a rather weak interaction^61^. We speculate that the binding of MK-886 to the PRR/MTBR of tau may shield these interactions and lead to an opening of the two termini, resulting in the observed decrease in FRET of the intramolecular FRET biosensor. From our previous observations with smFRET on tau constructs, this type of conformational change is often accompanied by an increase in FRET of the PRR/MTBR in recombinant protein systems^60^. In addition, in the *inter-molecular* FRET biosensor, MK-886 does not completely abolish the FRET signal, further suggesting that tau-tau interactions are not completely disrupted but may have undergone a conformational change from an oligomer with closer proximity between the fluorophores to one with a more open conformation.

The other four hit compounds identified from our HTS did not display direct binding to tau protein and thus are most likely acting through indirect mechanisms to attenuate tau oligomer-induced FRET. Although outside the scope of the present study, these potential indirect effectors of tau oligomerization may prove relevant as hit compounds, targeting orthogonal pathological phenotypes in AD and related tauopathies. For example, the indirect hit benzbromarone is an anti-hyperuricemia agent which attenuates oxidative stress, a known cellular dysfunction in AD^10^, and has been shown to have neuroprotective effects and reduce the risk for neurodegenerative diseases such as dementia^64^. Two additional indirect hit compounds, bumetanide and torsemide, are both loop diuretics which inhibit the sodium-potassium-chloride cotransporter (NKCC1) in vascular smooth muscle and have been shown to reduce the risk of AD dementia in both adults with normal cognition or with mild cognitive impairments^65^. Lastly, the fourth indirect hit compound, triclosan, is an antibacterial and antifungal agent but has been recently shown to induce autophagy, another cellular process that is commonly compromised in tauopathies^66^. The relevance of these compounds to tauopathies or AD supports the competency of identifying indirect hits as an added advantage of our cellular approach over purified protein assays.

The identity of a specific, toxic tau oligomeric species remains elusive. Indeed, it is unlikely that a single, unique toxic conformation exists. It is far more likely that an ensemble of toxic oligomers (differing in size, conformation and even molecular constituency) populates the amylogenic cascade^29–35^. This heterogeneity in potential tau oligomer targets highlights the need for an ultrasensitive screening platform capable of monitoring structural changes within the ensemble of tau assemblies. Our FRET-based platform for monitoring full-length tau oligomerization in cells is a new technology that is capable of elucidating novel compounds which alter conformation and oligomerization states of tau, thereby providing a new pipeline of therapeutic discovery for tauopathies. One aspect of our technology that will be straightforward to improve will be to express tau in a stable or inducible cell line^44^. This will further improve conditions for screening in a native environment. Furthermore, in a larger screen, the high information content of time-resolved FRET has the potential to resolve hit compounds into distinct classes based on their structural effects on the target^48^. This strategy, combining fluorescent biosensor and time-resolved FRET measurement in a HTS platform, should be generally applicable to other drug discovery efforts targeting intrinsically disordered proteins involved in numerous neurodegenerative diseases.

## Methods

### Molecular biology

To generate tau-GFP and tau-RFP, cDNA encoding full-length 2N4R tau (441 amino acids) was fused to the N-terminus of EGFP and TagRFP vectors. The P301L mutation was introduced by QuikChange mutagenesis (Agilent Technologies, Santa Clara, CA) and sequenced for confirmation. The GFP-tau-RFP was generated by fusing the N-terminal of tau to the C-terminus of GFP and the C-terminus of tau to the N-terminus of RFP. All constructs contain the monomeric mutation A206K to prevent constitutive fluorophore clustering^67^.

### Cell culture and generation of stable cell lines

HEK293 and SH-SY5Y cells (ATCC) were cultured in phenol red-free Dulbecco’s Modified Eagle Medium (DMEM, Gibco) supplemented with 2 mM L-Glutamine (Invitrogen), heat-inactivated 10% fetal bovine serum (FBS HI, Gibco), 100 U/ml penicillin and 100 μg/ml streptomycin (Gibco). Cell cultures were maintained in an incubator with 5% CO2 (Forma Series II Water Jacket CO2 Incubator, Thermo Scientific) at 37 °C. The *inter-molecular* tau FRET biosensor was generated by transiently transfecting HEK293 cells using Lipofectamine 3000 (Invitrogen) with tau-GFP and tau-RFP (1:20 DNA plasmid concentration ratio). The effectiveness of HEK293 cells transfected with FRET constructs as a HTS platform has been demonstrated in our previous work^47–48^. To generate stable cell lines expressing GFP-tau-RFP or tau-GFP only, HEK293 cells were transiently transfected using Lipofectamine 3000 with GFP-tau-RFP or tau-GFP DNA plasmids. Transiently transfected cells were treated with G418 (Enzo Life Sciences, Farmingdale, NY) to eliminate non-expressing cells. Stable cell lines expressing GFP-tau-RFP or tau-GFP with the largest population of expressing cells were selected by fluorescence microscopy. The GFP-linker-RFP (linker contains 32 amino acids, GFP-32AA-RFP) control stable cell line was generated as described previously^68^. The control cells expressing only free soluble fluorophores (GFP or RFP only) were generated by transiently transfecting HEK293 cells using Lipofectamine 3000 with plasmids containing GFP or RFP DNAs at the same plasmid concentration as the *inter*-molecular tau FRET biosensor.

### Pilot screening with NIH clinical collection (NCC) library

The NIH Clinical Collection (NCC) library, containing 727 compounds, was purchased from Evotec (Hamburg, Germany), formatted into 96-well mother plates using an FX liquid dispenser, and formatted across three 384-well plates at 50 nL (10 μM final concentration/well) using an Echo liquid dispenser. DMSO (matching %v/v) was loaded as in-plate no-compound negative controls to make a total of 960 wells. The 384-well flat, black-bottom polypropylene plates (PN 781209, Greiner Bio-One) were selected as the assay plates for their low autofluorescence and low interwell cross talk. The plates were sealed and stored at −20 °C until use. Two days prior to screening, HEK293 cells were transfected using Lipofectamine 3000 with tau-GFP/RFP (tau FRET biosensor) in 15 x 100 mm plates (5 x 10^6^ cells/plate) and the stable tau-GFP cell line (donor-only control) was expanded in five 225 cm^2^ flasks. On each day of screening, the compound plates were equilibrated to room temperature (25 °C). The cells were harvested from the 100 mm plates by incubating with TrypLE (Invitrogen) for 5 min, washed three times in PBS by centrifugation at 300 g and filtered using 70 μm cell strainers (BD Falcon). Cell viability, assessed using a trypan blue assay, was >95%. Cells were diluted to 1 million cells/ml using an automated cell counter (Countess, Invitrogen). Expression of tau-GFP and tau-GFP/RFP (tau FRET biosensor) was confirmed by fluorescence microscopy prior to each screen. After resuspension and dilution in PBS, the biosensor cells were constantly and gently stirred using a magnetic stir bar at room temperature, keeping the cells in suspension and evenly distributed to avoid clumping. During screening, cells (50 μl/well) were dispensed by a Multidrop Combi Reagent Dispenser (Thermo Fisher Scientific) into the 384-well assay plates containing the compounds and allowed to incubate at room temperature for 2 hours before readings were taken by the fluorescence lifetime plate reader (Fluorescence Innovations, Inc) as described previously^47–48^.

### HTS data analysis

As described previously^47–48^, time-resolved fluorescence waveforms for each well were fit with single-exponential decays using least-squares minimization global analysis software to give donor-acceptor lifetime (τ_DA_) and donor-only lifetime (τ_D_). FRET efficiency (*E*) was then calculated based on Equation 1.

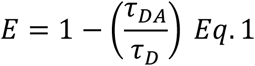

Assay quality was determined with the lead compound (MK-886) as positive control and DMSO as a negative control and calculated based on Equation 2^69^,

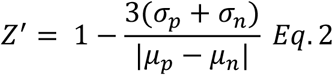

where σ_p_ and σ_n_ are the standard deviations (SD) of the observed τ_DA_ values, and μ_p_ and μ_n_ are the mean τ_DA_ values of the positive and negative controls. To make this metric less sensitive to strong outliers, we utilized the normalized median absolute deviation (1.4826*MAD) and median in place of the standard deviation and mean, respectively^70^.

Fluorescent compounds were flagged as potential false positives due to interference from compound fluorescence by a set of stringent fluorescent compound filters based on analysis of the spectral waveforms of each well from the NCC screen^47–48^. After removal of fluorescent compounds, a histogram of the FRET distribution from all compounds in the screen was plotted and fit to a Gaussian curve to obtain the mean (μ) and standard deviation (σ, SD). A hit was defined as a compound that decreased the FRET efficiency by more than five times the standard deviation (5σ) relative to the mean μ. Five reproducible hits, MK-886 (Cayman Chemical), Benzbromarone (Millipore Sigma), Bumetanide (Millipore Sigma), Torsemide (Millipore Sigma) and Triclosan (Millipore Sigma) were purchased.

### Protein purification

Full-length 2N4R tau proteins were purified from *E. coli* using previously published protocols^60^. Full-length tau was expressed with a cleavable His-tag. After elution from a nickel column, cleavage of the His-tag was achieved by incubation with tobacco etch virus (TEV) protease at room temperature for at least 4 hours, followed by passing through the His-tag column again to separate cleaved and uncleaved protein and remove the TEV. Final purification was performed by size-exclusion chromatography and the purity of the proteins was assessed by 4%-15% SDS-PAGE gels (Bio-Rad) under reducing conditions, followed by Coomassie staining. Fractions of pure proteins from the gels were pooled together and the protein stock concentrations were measured using the BCA assay (Thermo Fisher Scientific).

### Surface plasmon resonance (SPR) binding assay

Binding affinity between full-length WT 2N4R tau and the hit compounds was determined by SPR analysis using BIAcore S200. Recombinant tau proteins were immobilized on the CM5 sensor chip (Biacore, GE Healthcare) via amine coupling. Briefly, the dextran surface was activated with a 1:1 mixture of 0.4 M 1-ethyl-3-(3-dimethylaminopropyl)carbodiimide hydrochloride and 0.1 M N-hydroxysuccinimide. Tau protein (20 μg/ml) in 10 mM sodium acetate at pH 3.5 was flowed past a working surface before blocking the remaining activated carboxymethyl groups with 1 M ethanolamine at pH 8.5 to achieve a level of 1200 RU suitable for binding analysis. The reference surface was activated and reacted with only ethanolamine.

For direct binding assays to the tau protein, hit compounds at eight different concentrations (1 nM to 5 μM), as well as DMSO-only controls, were prepared in HEPES-EP containing a total of 2% DMSO. The samples were injected over both the reference and tau immobilized surfaces at 10 μl/min for 180 seconds and dissociated in glycine-HCl pH 2.5. All the samples, along with blanks from buffer and DMSO-only controls, were measured on a 96-well microplate (Biacore, GE Healthcare) at 25 °C. Reflectivity response data points were extracted from response curves at 5 seconds prior to the end of the injection to determine steady-state binding. All the data were double referenced with blanks using standard procedures with Biacore S200 Evaluation Software v1.0.

### FRET dose-response assay

MK-886, which shows direct binding to tau proteins and the strongest change in FRET, was tested in a FRET dose-response assay. The compound was dissolved in DMSO to make a 10 mM stock solution, which was serially diluted in 96-well mother plates. MK-886 was screened at nine different concentrations (1 nM to 10 μM). Compound (1 μl) was transferred from the mother plates into assay plates using a Mosquito HV liquid handler (TTP Labtech Ltd, UK). Three days prior to conducting the assays, the stable GFP-tau-RFP cells and GFP-32AA-RFP control cells were expanded in two 225 cm^2^ flasks (Corning). The preparations for tau-GFP/RFP FRET biosensors and the soluble GFP/RFP controls cells were carried out similar as above.

### Cell cytotoxicity assay

Cell cytotoxicity was measured using the CytoTox-Glo (Promega Corporation) luminescence assay kit. SH-SY5Y human neuroblastoma cells were plated at a density of 5 x 10^6^ cells/plate in 100 mm plate (Corning) and transfected with unlabeled full-length WT 2N4R tau or equivalent vector-only control for 24 hours. The transfected cells were then plated at a density of 5000 cells/well in white solid 96-well plate (Corning) with a total volume of 100 μl, followed by treatment with MK-886 at eight different concentrations (1 nM to 5 μM), as well as DMSO-only controls, for another 72 hours. After incubation, 50 μl of CytoTox-Glo Cytotoxicity Assay Reagent was added to all wells followed by mixing by orbital shaking and incubation for 15 minutes at room temperature. The first luminescence reading was measured using a Cytation3 Cell Imaging Multi-Mode Reader luminometer (BioTek). 50 μl of Lysis Reagent was then added, followed by incubation at room temperature for 15 min, and luminescence was measured again using the luminometer. Cell cytotoxicity was calculated by dividing the first luminescence signal by the second.

### Western blot analysis

To test the expression of tau FRET biosensors, HEK293 cells were plated in a six-well plate at a density of 1 x 10^6^ cells/well and transfected with tau-GFP/RFP (tau FRET biosensor) plasmid. To test the clearance and phosphorylation state of tau in the cytotoxicity assay, SH-SY5Y cells were plated in a six-well plate at a density of 1 x 10^6^ cells/well and transfected with unlabeled tau plasmid for 24 hours followed by treatment of MK-886 (5 μM) for 72 hours. In both cases, cells were lysed for 30 minutes on ice with radioimmunoprecipitation assay (RIPA) lysis buffer (Pierce RIPA buffer, Thermo Fisher Scientific) containing 1% protease inhibitor (Clontech, Mountain View, CA) and 1% phosphatase inhibitors (Millipore Sigma), and centrifuged at 15,000 g at 4 °C for 15 min. The total protein concentration of lysates was determined by bicinchoninic acid (BCA) assay (Pierce), and equal amounts of total protein (60 μg) were mixed with 4× Bio-Rad sample buffer and loaded onto 4%—15% Trisglycine sodium dodecyl sulfate-polyacrylamide gel electrophoresis (SDS-PAGE) gels (Bio-Rad, Hercules, CA). Proteins were transferred to polyvinylidene fluoride (PVDF) membrane (Immobilon-FL, EMD Millipore, Billerica, MA) and probed using antibodies against tau (Tau-5, Thermo Fisher Scientific) or antibody specific to Serine 396 of tau (Phospho-Tau S396, Thermo Fisher Scientific) with β-actin (ab8227, Abcam, Cambridge, MA) used as loading control. Blots were quantified on the Odyssey scanner (LI-COR Biosciences, Lincoln, NE).

### Protein labelling and single-molecule FRET (SmFRET) measurements

For site-specific labeling with maleimide-reactive fluorophores, cysteine residues were introduced using QuikChange Site-Directed Mutagenesis (Stratagene). Naturally occurring cysteines were mutated to serines. Labelling positions were selected to roughly mark the boundaries of the N-terminal domain or the proline-rich and microtubule-binding region of tau. Tau protein was purified as described above and labeled immediately following purification following published protocols^60^. Briefly, the protein (typically 200 μL of ~100 μM protein) was incubated with 1 mM DTT for 30 minutes at room temperature followed by exchange into labeling buffer (20 mM Tris pH 7.4, 50 mM NaCl, and 6 M guanidine HCl) to remove DTT. The protein was incubated with the donor fluorophore, Alexa Fluor 488 maleimide (Invitrogen), at a protein to dye ratio of 2:1 at room temperature for one hour with stirring. The acceptor dye, Alexa Fluor 594 maleimide (Invitrogen), was added at a 5-fold molar excess and incubated overnight at 4°C with stirring. Excess dye was then removed by buffer-exchanging the labeled solution into 20 mM Tris (pH 7.4) and 50 mM NaCl buffer using Amicon concentrators (Millipore) and then passed over two coupled HiTrap Desalting Columns (GE Life Sciences).

Single-molecule FRET measurements were carried out using ~30 pM of labelled tau in phosphate buffer (40 mM potassium phosphate, 50 mM KCl, pH 7.4) in 8-chambered Nunc coverslips (ThermoFisher) passivated with poly(ethylene glycol) poly(L-lysine) (PEG-PLL) to reduce protein adsorption to the chambers. Control measurements included DMSO to match the concentration in samples containing MK-886. Measurements were made on a MicroTime 200 time-resolved confocal microscope (Picoquant) in pulsed interleaved excitation FRET (PIE-FRET) mode. Laser power from 485 and 561 nm lasers, operated at 40 MHz pulse rate, was adjusted to ~30 μW before sample illumination. Fluorescence emission was collected through the objective and passed through a 150 μm diameter pinhole. Photons were separated by an HQ585LP dichroic in combination with ET525/50M and HQ600LP filters and detected by avalanche photodiodes. Photon traces were collected in 1 ms time bins for one hour. A cutoff of 25 counts/ms was applied to discriminate between bursts arising from fluorescently labeled protein and background noise. No bursts were identified in photon traces with DMSO only and MK-886 only when this criterion was applied. The FRET efficiency (ETeff) was calculated using SymphoTime 64 software. SmFRET histograms were fit with Gaussian distributions to determine the peak ETeff values. Alignment of instrument and analysis were verified using 10 base pair, 14 base pair and 18 base pair dsDNA standards.

### Thioflavin-S (ThS) assay

HEK293 cells were transfected with tau-RFP (at equivalent DNA concentration as used in the tau FRET biosensor) for 48 hours prior to the addition of preformed fibrils (PFF). Tau-GFP was not used, as it would interfere with the Thioflavin-S (ThS) signal. To make the PFF, 100 μl of purified tau proteins (10 μM) with DTT (5 mM) and heparin (0.4mg/ml) were first incubated for 120 hours at 37°C and shook at 1000 rpm in a thermal shaker (Thermo Fisher Scientific). After incubation, the sample was subjected to ultracentrifugation at 80,000 rpm for 30 minutes. The pellet was collected and sonicated to break up the fibrils into smaller pieces. The concentration of the fibrils was then measured by BCA. The sonicated fibrils were then treated to the transfected cells at a concentration of 40 μg/ml for 24 hours before conducting the ThS assay. Thioflavin-S (ThS, Millipore Sigma, product no. T1892) was dissolved in PBS buffer and was filtered through a 0.2 μm syringe filter to make a stock solution of 2.5 mM. For the ThS assay, cells were fixed with 1 ml of 4% paraformaldehyde in TBS for 15 minutes followed by washing with 1 ml of TBS for 5 minutes twice. After fixing, cells were permeabilized with 1ml of 1% Triton in TBS for 5 minutes, followed by washing with 1ml of TBS for 5 minutes twice. After permeabilization, cells were then treated with 0.002% ThS in TBS and incubate in the dark for 20 minutes. Cells were then washed twice with 50% ethanol for 10 minutes each and finally washed twice with TBS for 5 minutes each. Cells were then imaged with a fluorescence microscope using EVOS-FL cell imaging systems at 20X magnification.

### Thioflavin-T (ThT) assay

Thioflavin-T (ThT, Sigma, product no. T3516) was dissolved in PBS buffer and was filtered through a 0.2 μm syringe filter to make a stock solution of 2.5 mM. ThT was then diluted to 20 μM prior to addition to the tau proteins. The samples for ThT measurements were prepared by mixing 25 μl of 20 μM tau proteins with 25 μl of 20 μM of ThT, resulting in final concentrations of 10 μM tau proteins and 10 μM ThT. DTT (5 mM) and heparin (0.4 mg/ml) were then added to the samples; a control sample lacked addition of heparin. Lastly, the samples were treated with MK-886 (0.5 μM or 5 μM) and gossypetin (50 μM) with DMSO added to the no-compound controls. The ThT samples (50 μl each) were transferred to a black 96-well non-binding surface microplate with clear bottom (Corning product no. 3655) and incubated at 37°C with mild shaking (200 rpm) in the Cytation 3 plate reader. The ThT fluorescence was measured by the Cytation 3 plate reader through the bottom of the plate with excitation filter of 440 nm and emission filter of 480 nm. Readings were acquired every 20 minutes for a total of 120 hours.

### Statistical analysis

Data are shown as mean ± standard deviation unless stated otherwise. Statistical analysis was conducted by an unpaired Student’s t-test using GraphPad Software to determine statistical significance for all experiments. Values of P <0.05 were considered statistically significant.

## Acknowledgements

We thank Nagamani Vunnam, Malaney C. Young and Breeanne M. Brand from the Sachs Group, Tory Schaaf, Samantha Yuen, Andrew Thompson, and Razvan Cornea from the Thomas Group and Benjamin Grant from Fluorescence Innovations, for technical support and discussions. We also thank Dr. David Odde from University of Minnesota for discussions. Compound dispensing and surface plasmon resonance (SPR) at the UMN Institute of Therapeutic Drug Discovery and Development (ITDD) High-Throughput Screening Laboratory, and spectroscopy at the UMN Biophysical Technology Center. This research uses technology patented by the University of Minnesota, with an exclusive commercial license to Photonic Pharma LLC. The authors disclosed receipt of the following financial support for the research, authorship, and/or publication of this article: This study was supported by U.S. National Institutes of Health (NIH) grants to J.N.S. (R01AG053951) and D.D.T. (R37AG026160). C.H.L. was supported by a Doctoral Dissertation Fellowship from the University of Minnesota.

## Author contributions

C. H.L. designed and conducted the experiments. C.K.W.L. and Z.D. contributed to protein purification and provided assistance to cell-based assays and biochemical experiments. S.W. and E. R. performed and analyzed the single-molecule FRET experiment. D.D.T. provided expertise on FRET and HTS, and provided comments and edits to the manuscript. A.R.B. provided comments and edits to the manuscript. C.H.L and J.N.S. wrote the manuscript.

## Materials & Correspondence

Correspondence and material requests should be sent to Jonathan Sachs.

---

**Supplementary Fig. 1:**
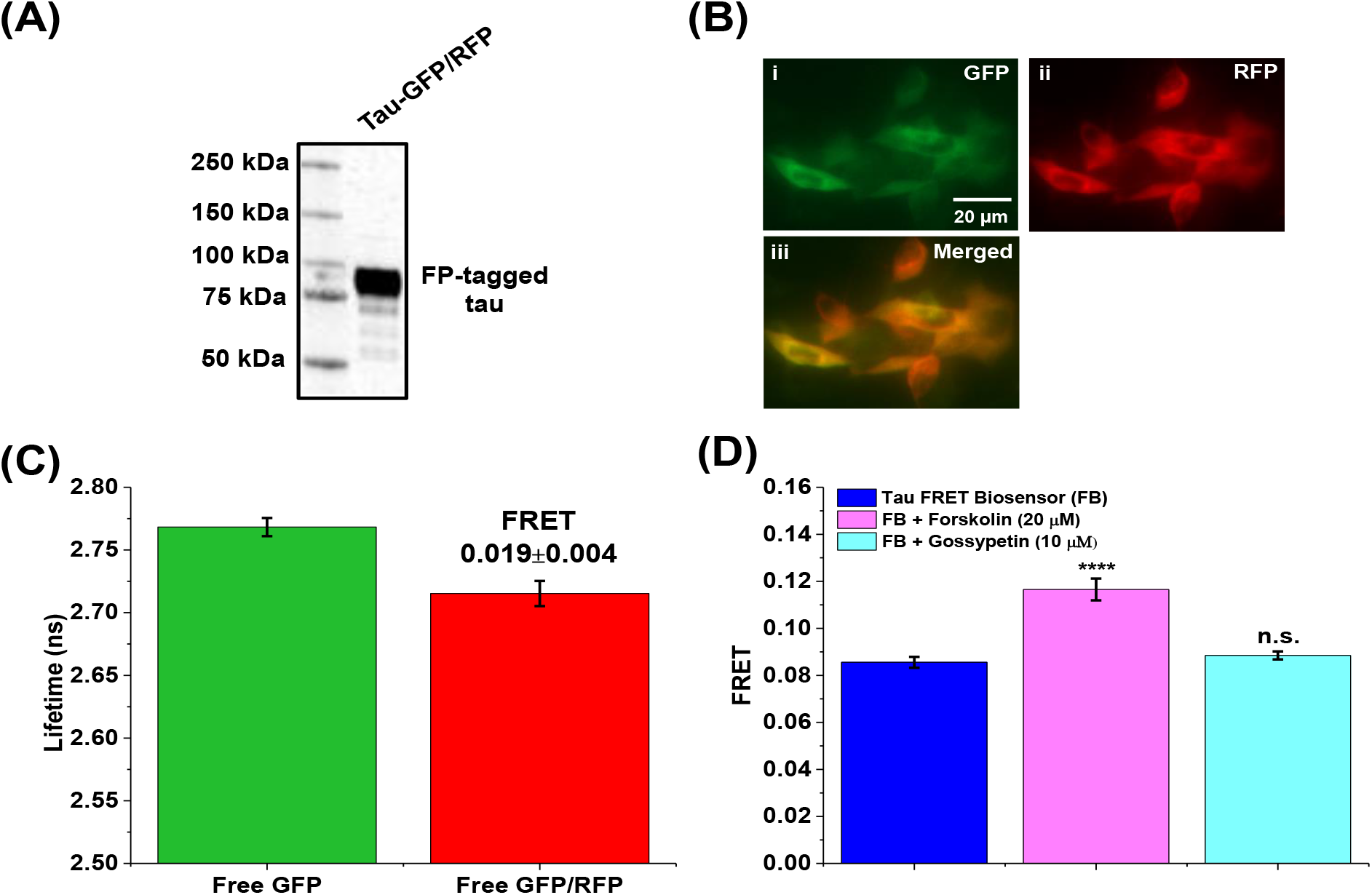
Characterization of tau *inter*-molecular FRET biosensor and soluble free GFP/RFP-only expressed in HEK293 cells as controls. (A) Expression of tau-GFP and tau-RFP expressed in HEK293 cells stained with Tau-5 antibody. (B) Fluorescent microscopy images of soluble free GFP/RFP-only expressed in HEK293 cells at the same donor-to-acceptor ratio as the biosensor show the co-expression and co-localization of the fluorophores in the cells. (C) Fluorescence lifetime measurements of the GFP/RFP-only controls show a FRET of 0.019±0.004, indicating the basal FRET from free soluble fluorophore. (D) Functionality of the tau FRET biosensor was confirmed by addition of forskolin (a known small molecule that induces tau hyperphosphorylation and self-association) and gossypetin (a known inhibitor of fibril formation). N=3, ****p<0.0001 and n.s. indicates not significant.

**Supplementary Fig. 2:**
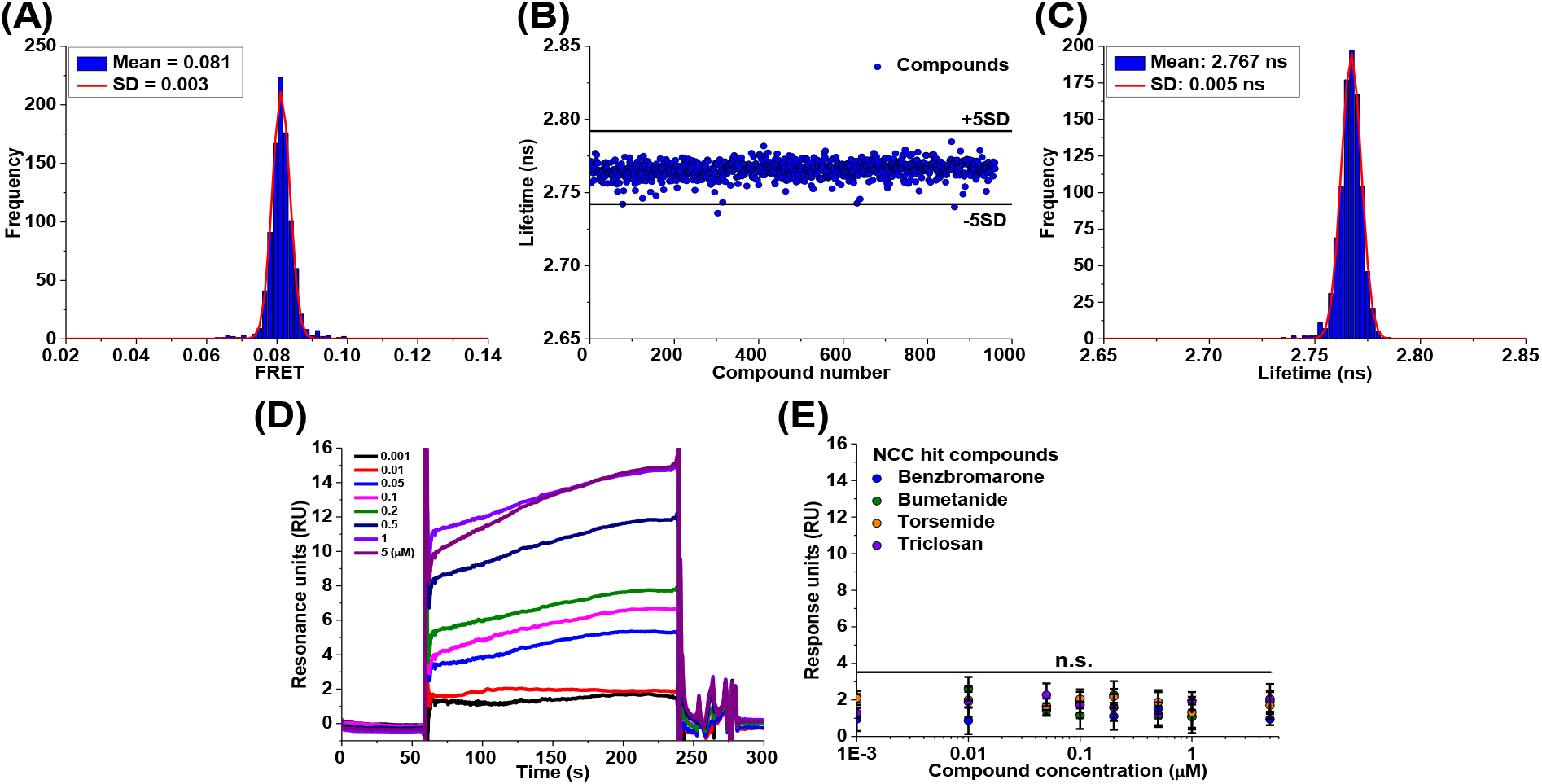
Donor-only control screen and surface plasmon resonance (SPR) characterization of hit compounds in binding to purified tau protein. (A) Histogram plots of all compounds from the NCC screen after removal of fluorescent compounds show an average FRET efficiency of 0.081 and a standard deviation (SD) of 0.003. (B) Representative donor-only control screen with NCC library using cells expressing only tau-GFP which do not show FRET signal so the lifetime plot is shown. Applied threshold at a change in lifetime of 0.025 (5SD) is shown by the black lines. There is no reproducible hit from the donor-screen indicating that the hits observed are due to random occurrence. In addition, the hit compounds obtained from the FRET screen do not appear as hits in any of the donor-only control screens. (C) Histogram plots of all compounds from the NCC donor-only screen after removal of fluorescent compounds show an average lifetime of 2.767±0.005 ns. (D) Raw curves of SPR characterization of binding between purified tau protein and MK-886. (E) SPR characterization of other four hit compounds shows no direct binding or interaction between the compounds and the tau protein. N=3 and n.s. indicates not significant.

**Supplementary Fig. 3:**
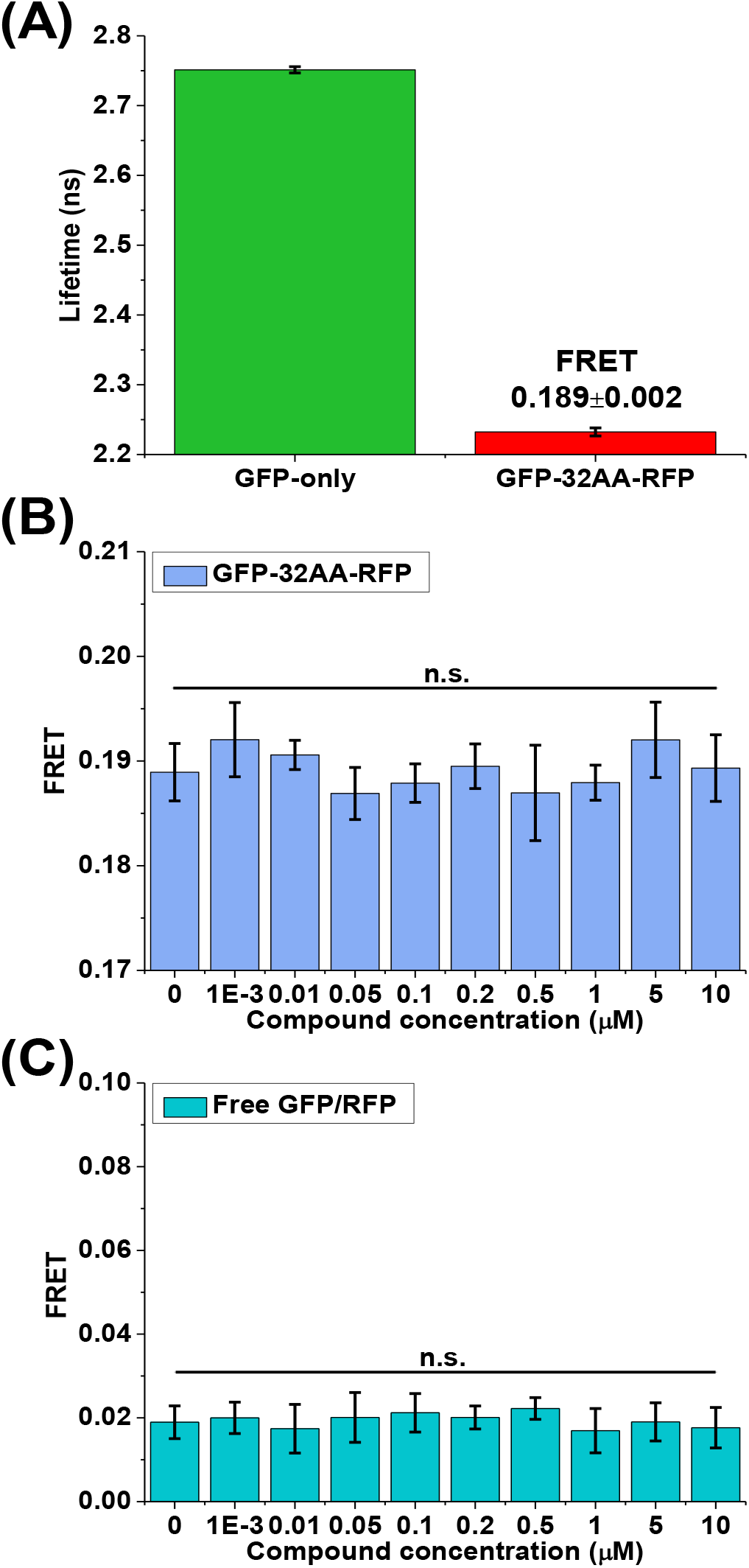
Characterization of MK-886 with control FRET biosensors. (A) The GFP-32AA-RFP expressing cells show a FRET of 0.189±0.002, which can be used as a control to test compound interference with GFP or RFP fluorophores, hence causing a FRET change in the screen. (B) MK-886 does not cause any significant FRET change in the GFP-32AA-RFP expressing control cells. (C) MK-886 does not cause any significant FRET change in the soluble free GFP/RFP expressing control cells. N=3 and n.s. indicates not significant.

**Supplementary Fig. 4:**
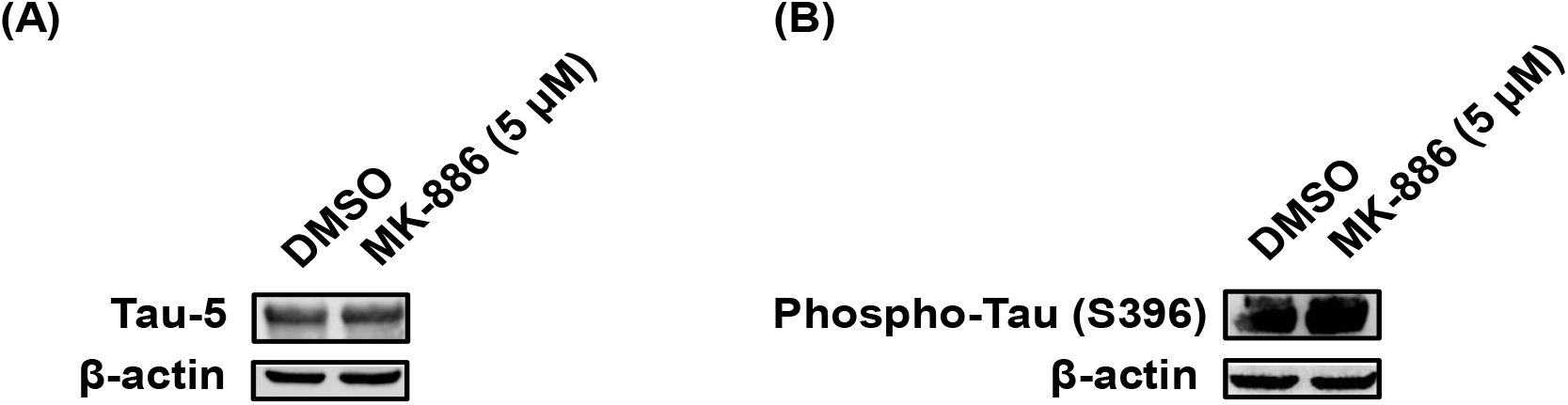
Controls for cell cytotoxicity functional assay in SH-SY5Y cells. (A) Total amount of tau expressed is shown by the Tau-5 antibody staining. MK-886 does not change the relative levels of expressed P301L tau. (B) The phosphorylation state Serine 396 of P301L tau expressed in SH-SY5Y cells is shown by the Phospho-Tau S396 antibody. The phosphorylation state of Serine 396 of P301L tau is not altered by MK-886. β-actin was used as loading control (N=3). Both controls indicate that the rescue of cell death is not due to an indirect mechanism.

**Supplementary Fig. 5:**
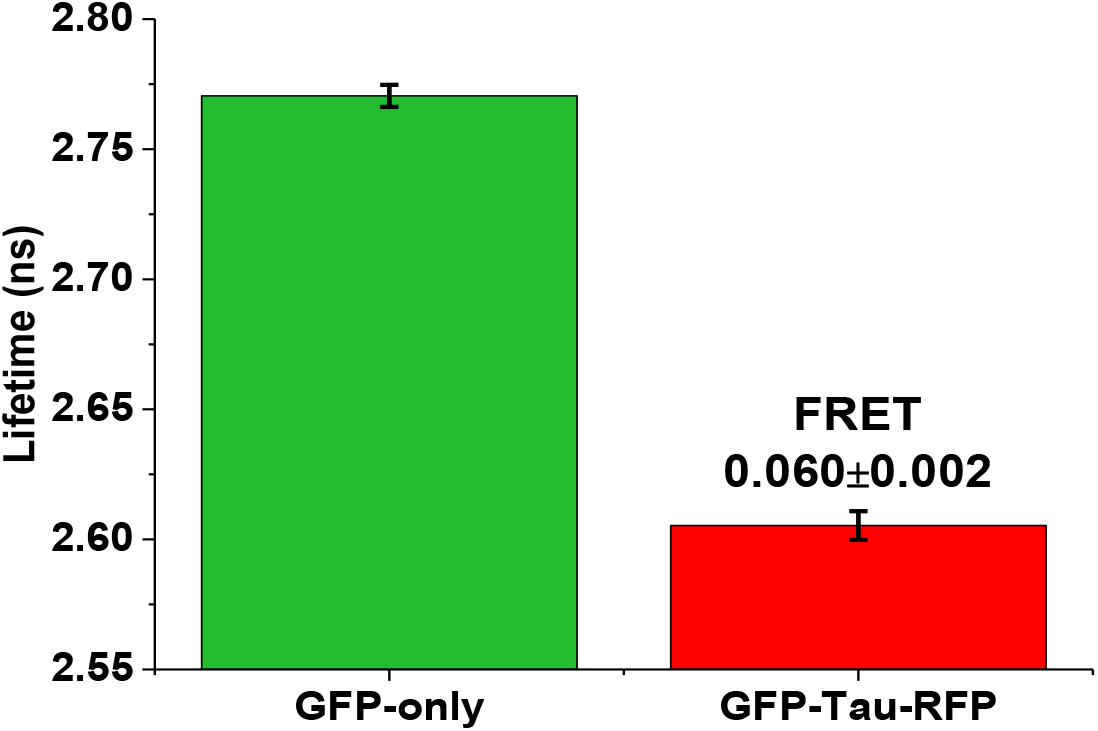
Lifetime measurements of the basal FRET of the cellular tau intrα-molecular biosensor. The cellular tau *intra*-molecular FRET biosensor shows a basal FRET of 0.060±0.002, illustrating the *intra*-molecular interactions arising from the paper-clip monomeric structure in which the N and C terminus of tau folded to close proximity of each other (N=3).

**Supplementary Fig. 6:**
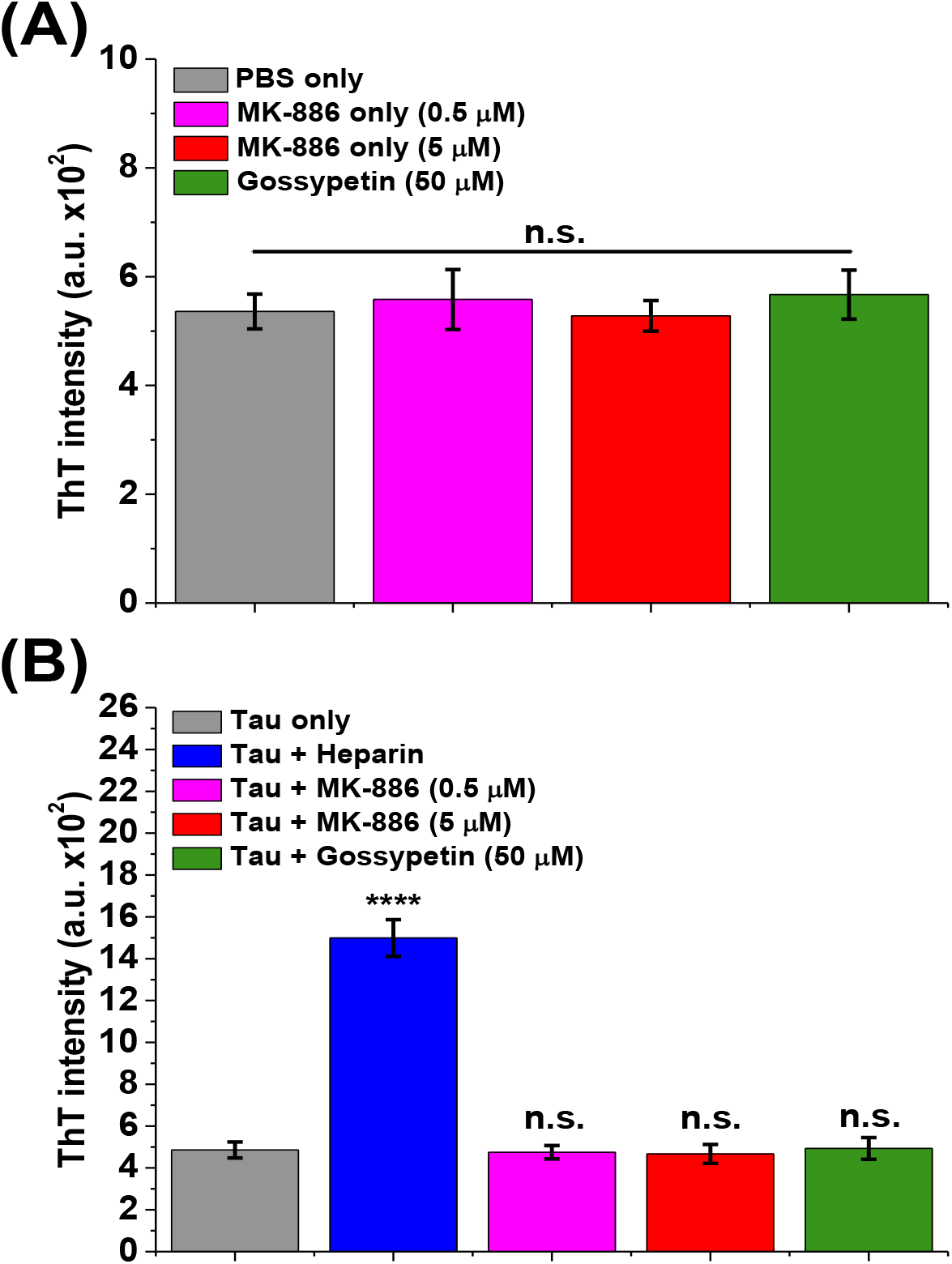
Controls for thioflavin-T (ThT) assay. (A) MK-886 and gossypetin do not interfere with ThT fluorescence after 72 hours of incubation. (B) MK-886 does not act as a nucleation center for fibril formation after 72 hours of incubation. Only the positive control of heparin shows a significant ThT positive signal. N=3, ****p<0.0001 and n.s. indicates not significant.

